# Characterisation of an antimicrobial and phytotoxic ribonuclease secreted by the fungal wheat pathogen *Zymoseptoria tritici*

**DOI:** 10.1101/130393

**Authors:** Graeme J. Kettles, Carlos Bayon, Caroline A. Sparks, Gail Canning, Kostya Kanyuka, Jason J. Rudd

## Abstract

- The fungus *Zymoseptoria tritici* is the causal agent of Septoria Tritici Blotch (STB) disease of wheat leaves. *Z. tritici* secretes many functionally uncharacterised effector proteins during infection. Here we characterised a secreted ribonuclease (*Zt6*) with an unusual biphasic expression pattern.
- Transient expression systems were used to characterise Zt6, and mutants thereof, in both host and non-host plants. Cell-free protein expression systems monitored impact of Zt6 protein on functional ribosomes, and *in vitro* assays of cells treated with recombinant Zt6 determined toxicity against bacteria, yeasts and filamentous fungi.
- We demonstrated that Zt6 is a functional ribonuclease and that phytotoxicity is dependent on both the presence of a 22-amino acid N-terminal “loop” region and its catalytic activity. Zt6 selectively cleaves both plant and animal rRNA species, and is toxic to wheat, tobacco, bacterial and yeast cells but not to *Z. tritici* itself.
- Zt6 is the first *Z. tritici* effector demonstrated to have a likely dual functionality. The expression pattern of Zt6 and potent toxicity towards microorganisms suggests that whilst it may contribute to the execution of wheat cell death, it is also likely to have an important secondary function in antimicrobial competition and niche protection.

## Introduction

In order to become a successful plant pathogen or parasite, micro-organisms must not only evolve strategies to manipulate host defences and acquire nutrients, but they must also interact with numerous other potential microbial competitors or antagonists present within their environmental niche. Control over these latter interactions are likely to be important for a pathogenic microbe to benefit fully from its own efforts to parasitize, and induce disease symptoms, on host plants.

For bacterial, fungal, viral and invertebrate pathogens there exists considerable knowledge of how they manipulate their plant hosts. Study of effectors from many different plant pathogens has revealed the important roles played by these secreted proteins in modifying host physiology and immunocompetence (Win *et al.*, 2012; Giraldo & Valent, 2013). The mechanisms by which pathogens resist host counter-defensive measures, and interact with other environmental microbes are lesser explored by comparison. How plant pathogenic microorganisms interact with other microbes during infection has often been studied in the area of biological control, but the molecular interplay underpinning these, often antagonistic, interactions are largely unknown.

The dothideomycete fungus *Zymoseptoria tritici* is a host- and tissue-specific pathogen of wheat leaves. Following spore germination on the leaf surface, there is a long phase of symptomless infection (typically at least six to seven days) whereby the fungus invades leaves through stomata and then colonises the mesophyll tissue (Kema *et al.*, 1996). During this phase, there is minimal activation of host defences. Shortly afterwards, there is a transition to the necrotrophic phase of the life cycle. This is accompanied by massive host defence reactions, host programmed cell death, release of nutrients and development of asexual fungal reproductive structures (pycnidia)(Kema *et al.*, 1996; Keon *et al.*, 2007).

The effector repertoire of *Z. tritici* has been partially characterised (Kettles & Kanyuka, 2016). Numerous bioinformatic and transcriptomic analyses have attempted to define the *Z. tritici* secretome through the infection cycle using several *Z. tritici* isolates (Mirzadi Gohari *et al.*, 2015; Rudd *et al.*, 2015; Palma-Guerrero *et al.*, 2016; Palma-Guerrero *et al.*, 2017). However, to date only a small number of candidate effector proteins have been functionally characterised. A Lysin domain effector (LysM effector) Zt3LysM (previously Mg3LysM), has been demonstrated to have chitin-binding properties (Marshall *et al.*, 2011) and is a functional ortholog of the *Cladosporium fulvum* Ecp6 effector (Bolton *et al.*, 2008; de Jonge *et al.*, 2010; Sánchez-Vallet *et al.*, 2013). The *Z. tritic*i *ΔMg3LysM* mutant is severely compromised in virulence (Marshall *et al.*, 2011) and this effector plays a crucial role in minimising host defence induction during early colonisation through suppression of chitin-triggered immunity (Lee *et al.*, 2014). The Necrosis and Ethylene-inducing Peptide 1 (NEP1)-like (NLP) family are widespread amongst plant pathogens. *Z. tritici* produces a single NLP effector (MgNLP) (Motteram *et al.*, 2009) which induces cell death in Arabidopsis but neither induces defence genes nor cell death in wheat. Recently, it was found that numerous *Z. tritici* effectors are recognised in the non-host tobacco species *Nicotiana benthamiana* (Kettles et al., 2017) and that recognition of several of these likely occurs at the plasma membrane-apoplast interface. Recently, the first avirulence gene *AvrStb6* which also encodes a small cysteine-rich effector, likely recognised by a wheat plasma membrane receptor-like protein was also recently described (Zhong *et al.*, 2017). However, in all cases to date, these examples of *Z. tritici* effector proteins all have roles only associated with the direct or indirect manipulation of plant defences in host or non-host plants.

In contrast to the examples above, a new concept of effector multi-functionality has recently gained traction based on data from other systems. The effector SnTox1 from the necrotrophic wheat pathogen *Parastagonospora nodorum* has recently been demonstrated to have a dual-functionality during wheat leaf infection (Liu *et al.*, 2016). Firstly, it induces a cell death pathway on wheat cultivars harbouring the *Snn1* sensitivity gene (Liu *et al.*, 2012; Shi *et al.*, 2016) and can then subsequently protect fungal hyphae from digestion by plant chitinases (Liu *et al.*, 2016) presumably activated as a consequence of the wheat cell death reaction. It has been speculated that pathogen effectors may have additional roles aside from direct host interactions, specifically regarding manipulation of local microbial communities (microbiomes) (Rovenich *et al.*, 2014). However, to date there are no published examples of single effector proteins with the capability to impact on the host as well as on host-associated microbial communities.

Secreted ribonucleases (sRNases) fulfil numerous roles in many different aspects of host-pathogen interactions both in animal and plant systems. Defensive RNases are secreted by human skin cells and their inhibition is essential for protection of the producing cells (Harder & Schröder, 2002; Thomas *et al.*, 2016). Two classes of sRNases of particular note are the highly toxic ribosome-interacting proteins (RIPs) and fungal ribotoxins (Lacadena *et al.*, 2007; Walsh *et al.*, 2013). The toxicity of these RNases is primarily due to their ability to cross cell membranes and to interact with ribosomes. They specifically attack the sarcin-ricin loop (SRL) of the larger eukaryotic rRNA, irreversible blocking protein synthesis and ultimately leading to cell death. For the fungal ribotoxins, much progress has been made at elucidating the modifications that confer these proteins potent toxicity in comparison to non-toxic RNases (Olombrada *et al.*, 2017). In plant-pathogen interactions, it has recently been demonstrated that several RNase-like effectors are secreted by the barley powdery mildew fungus *Blumeria graminis* f. sp. *hordei*. Whilst these secreted proteins are not functional RNases, they do significantly contribute to the disease progression of this biotrophic pathogen (Pedersen *et al.*, 2012; Pliego *et al.*, 2013).

All studies into the role of *Z. tritici* effectors to date have focussed on their function in wheat colonisation or potential recognition in non-host plants, whereas the potential for multi-functional effectors with roles in microbe-microbe interactions have not yet been explored. Here we describe a secreted ribonuclease (Zt6) from *Z. tritici* which is expressed specifically during distinct phases of wheat leaf infection and which possesses highly potent cytotoxic activity against plants as well as against various prokaryotic and eukaryotic microbes. This activity appeared to be reliant upon an N-terminal region of the protein which may facilitate cellular uptake into target cells. Conversely, Zt6 was found to be non-toxic to *Z. tritici* itself, suggesting the existence of an unknown self-protection mechanism.

## Materials and Methods

### Plant growth conditions

Seeds of *Nicotiana benthamiana* (tobacco) were sown and transplanted as described previously (Kettles *et al.*, 2017) and maintained in a glasshouse at 22°C/18°C (day/night) with a 16 h day-length. Seeds of *Triticum aestivum* (wheat) cultivar Riband were sown directly in half-trays of Rothamsted Prescription Mix and maintained in a glasshouse at a constant 17°C with a 16 h day-length.

### Fungal growth and inoculations

All *Z. tritici* strains were grown on YPD agar plates at 16°C for 5 days. Conidiospores were harvested in distilled H_2_O containing 0.01% Tween-20 and plant inoculations performed as described previously (Kettles *et al.*, 2017).

### Agrobacterium-mediated targeted deletion of *Zt6*

*Agrobacterium tumefaciens*-mediated transformation of the *Z. tritici* IPO323 *Δku70* strain (Bowler *et al.*, 2010) was performed as described previously (Motteram *et al.*, 2009). A construct designed to replace the *Zt6* (*Mycgr3G38105*) gene with the Hygromycin resistance gene was produced using the vector pCHYG and validated by diagnostic PCR. All primers used in production of the gene deletion construct and in sequence validation are available in Table S1.

### Agrobacterium-mediated transient expression of *Zt6* in *N. benthamiana*

Gateway cloning of *Zt6* into the binary vectors pEAQ-HT-DEST3 and pEARLEYGATE101 was conducted as described for other *Z. tritici* candidate effectors (Kettles *et al.*, 2017).

### Phylogenetic Analysis

Mature Zt6 amino acid sequence (lacking secretion signal peptide) was used as input for blastp analysis against all Dothideomycete sequences at NCBI (National Center for Biotechnology Information). The twenty closest homologs (Table S2) were aligned to Zt6 using ClustalW, and a phylogenetic tree constructed using the Maximum Likelihood (ML) method with 100 bootstrap replicates in MEGA6. Amino acid sequence representations of the predicted N-terminal “loop” and downstream ribonuclease (RNase) domains were produced using WebLogo 3 (http://weblogo.threeplusone.com) using the same ClustalW alignment as input.

### Site-directed mutagenesis

Generation of the *Zt6-H70A* mutant was performed using the QuikChange Lightning Multi Site Mutagenesis Kit (Agilent Technologies) following the manufacturer’s instructions. Primers were designed using the QuikChange Primer Design program and are included in Table S1.

### qRT-PCR

Expression analysis of Zt6 was performed using sample material generated from *Z. tritici* IPO323 grown in YPD media and at 1 day post-inoculation (dpi) of susceptible wheat cv. Riband. Total RNA was recovered using Trizol (Thermo Fisher Scientific) with a double chloroform extraction step. RNA preparations were treated with RQ1 DNase (Promega) following the manufacturer’s instructions. DNA-free RNA was recovered by ethanol precipitation, and 2 μg RNA was used as template for cDNA synthesis using SuperScript III reverse transcriptase (Thermo Fisher Scientific) and an oligo(dT)_20_ primer. cDNA was diluted 1:2 with distilled H_2_O and 1 μl of diluted cDNA was used in each reaction with SYBR Green Jumpstart Taq ReadyMix (Sigma). Triplicate reactions were prepared for each sample-primer pair combination and all reactions were performed using the following thermocycle on a Biorad CFX384 real-time system (3 min at 95°C, followed by 40 cycles of 30 s at 95°C, 30 s at 60°C, 30 s at 72°C, followed by melt curve analysis of 10 s at 95°C, then 65°C - 95°C in 0.5°C increments, 5 s at each). Relative expression values were calculated using the 2^-ΔCt^ method with *Z. tritici* β-tubulin as the reference gene. Expression values were rescaled for presentation such that the YPD treatment for each effector is equal to 1. All primer sequences are provided in Table S1.

### Wheat leaf sheath biolistic bombardment

*Zt6* variants were cloned into the pRRes14_RR.1m201_125 biolistic bombardment vector linearised by double-digestion with *Nco*I and *Spe*I restriction endonucleases. Corresponding *Nco*I/*Spe*I recognition sites were added to *Zt6* fragments by PCR amplification followed by double-digestion with the corresponding restriction endonucleases. *Zt6* fragments were incorporated into linearised pRRes14_RR.1m201_125 using T4 DNA ligase and transformed into One Shot TOP10 electrocompetent *E. coli* (Thermo Fisher Scientific) using standard procedures. Constructs were sequence verified by Sanger sequencing. All primer sequences used are available in Table S1.

For bombardments, young tillers were collected from ∼4-week old wheat cv. Apogee plants. A section ∼10 cm long containing the immature inflorescence was cut from each tiller and the ends sealed with parafilm. The trimmed stems were surface sterilised using 70% v/v ethanol for 3 mins and 10% v/v domestic thin bleach (sodium hypochlorite content of 4-6%) for 3 mins followed by several repeat washes with sterile water. Leaf sheaths were isolated by cutting away the outer layers to expose the immature inflorescence, and approximately 1-1.5cm of the young leaf sheath closest to the inflorescence was removed. The isolated leaf sheaths were plated on L7 medium supplemented with 3% w/v sucrose, 0.5 mg/l 2,4-D (Sigma-Aldrich, UK) and 10 mg/l AgNO_3_ (Sigma-Aldrich, UK), solidified with 5 g/l agargel (Sigma-Aldrich, UK) in 9 cm petri dishes, placing sufficient leaf sheath sections to cover the central 2 cm^2^. The prepared leaf sheaths were transformed on the same day as isolation. Particle bombardment was carried out according to (Sparks & Jones, 2014). Test constructs were precipitated onto 0.6 μm gold particles (Bio-Rad, UK) alongside a construct which contained the green fluorescent protein (GFP) reporter gene under control of the maize ubiquitin gene promoter with an *Arabidopsis thaliana* histone H2B-like nuclear targeting sequence and nopaline synthase gene (nos) terminator. The particles were delivered into target tissues using the Bio-Rad PDS-1000/HeTM particle gun with a rupture pressure of 650 psi and a vacuum of 28-29” Hg. Treated leaf sections were incubated at ∼22°C in the light (12 hr photoperiod) for 2 days prior to visualisation of the GFP using a Leica M205 FA stereomicroscope with a fluorescence filter suitable for GFP.

### RNase activity assays

Wild type (FL) and mutant Zt6 (Δ19-40, H70A, Δ19-40+H70A) were recombined into the Gateway-compatible pDEST17 expression vector (Invitrogen) using LR clonase II enzyme mix following the manufacturer’s instructions. Sequence-verified constructs were linearised by digestion with *EcoRV* restriction endonuclease and 1 μg linearised plasmids were used as template for *in vitro* transcription reactions (carried out for 3 h at 37°C) using the MEGAscript T7 transcription kit (Ambion). Transcription products were purified using the RNA cleanup protocol of the RNeasy Mini Kit (Qiagen). *In vitro* translation reactions were assembled using the Rabbit Reticulocyte Lysate/Wheat Germ Extract Combination System (Promega) using 1 μg RNA as template. The provided Luciferase control RNA was used as negative control for RNase activity. Reactions were incubated for 10-90 minutes at 30°C (rabbit reticulocyte lysate) or 25°C (wheat germ extract). Ribosomal RNA was recovered following the method of (Kao *et al.*, 2001) before resolving on 2% semi-denaturing agarose gels.

### *Tobacco rattle virus* (TRV)-mediated Virus-induced gene silencing (VIGS) in *N. benthamiana*

This method has been fully described previously (Kettles *et al.*, 2017). Briefly, 2-3 week-old *N. benthamiana* seedlings were Agroinfiltrated with a 1:1 mix of Agrobacterium strains harbouring PTV00 (TRV RNA2)-derived constructs and Agrobacterium GV3101 carrying pBINTRA6 (TRV RNA1) at a final OD_600_ = 1 to initiate silencing. After 2-3 weeks, full-length Zt6 was expressed in systemically-infected leaves by Agroinfiltration of GV3101, carrying pEAQ-HT-DEST3 Zt6, at OD_600_=1.2. Leaves were visually assessed for cell death symptoms at 7 dpi.

### Recombinant protein expression and purification

Zt6 variants were cloned into the p75 protein expression vector (Franco-Orozco *et al.*, 2017) linearised by digestion with *Pac*I using yeast recombination cloning in *Saccharomyces cerevisiae* FY834. Recombined plasmid DNA (p75 Zt6 Δ19-40) or (p75 Zt6 H70A) was recovered from *S. cerevisiae* and bulked up by transformation into One Shot TOP10 electrocompetent *E. coli* (Thermo Fisher Scientific). Plasmids were sequence-verified by Sanger sequencing and constructs were linearised by digestion with *Pme*I prior to electrotransformation into *Pichia pastoris* GS115 cells. *P. pastoris* (p75 Zt6 Δ19-40) or (p75 Zt6 H70A) transformants were selected by growth on YPD supplemented with zeocin (100 μg ml^-1^) for 2-3 days at 30°C. For identification of colonies highly expressing recombinant proteins, 2 ml YPD cultures were prepared for 10-15 individual colonies of each construct and grown for 48 h at 30°C (250 rpm) in a shaker incubator. Cells were pelleted by centrifugation (2000 x g for 5 min) and 2 μl supernatant spotted on nitrocellulose membrane (Amersham Protran Premium 0.45 NC, GE Healthcare). Membranes were incubated in blocking buffer (TBST + 2% milk powder) and then probed with anti-V5 HRP conjugated monoclonal antibody (E10/V4RR, Pierce) at 1:5000 dilution in blocking buffer before washing in TBST. Signal was detected using Amersham ECL prime kit (GE Healthcare) and exposure to X-ray film. For scaled-up purification, transformants highly expressing recombinant proteins were grown flasks containing 1.5 L YPD supplemented with zeocin (50 μg ml^-1^) for 48 h at 30°C (250 rpm) in a shaker incubator. Cells were pelleted by centrifugation, and supernatant incubated with anti-V5 affinity gel (Biotool/Stratech Scientific Ltd.) following the manufacturer’s instructions. Affinity gel was washed thrice in TBS and protein recovered by 5 x 1 ml elutions with elution buffer (0.25x TBS, 200 μg ml^-1^ V5 peptide). Protein elutions were pooled, filter sterilised and buffer-exchanged to distilled H_2_O using Amicon Ultra-15 columns (3KDa MWCO, Merck Millipore) such that final concentration of V5 peptide was ≤ 1 μg ml^-1^. Purified proteins were quantified by spectrophotometry measuring absorbance at 280 nm (A_280_).

### Microbial toxicity assays

LB or YPD liquid cultures with appropriate antibiotics were inoculated with *E. coli* Top10 (pEAQ-HT-DEST3 Avr4), *P. pastoris* GS115 (p75 EV) or *S. cerevisiae* FY834 and grown overnight at 30°C. Cells were pelleted by centrifugation at 4000 rpm for 5 mins, washed once in distilled H_2_O and recovered as above. *E. coli* cells were diluted with distilled H_2_O to a final OD_600_ = 0.01, with *P. pastoris* and *S. cerevisiae* diluted to OD_600_ = 0.6. The *Z. tritici* IPO323 *Δku70* isolate was grown for 5 days at 16°C on YPD agar plates supplemented with Geneticin. Conidiospores were resuspended to a density of 1x10^7^ spores/ml in distilled H_2_O + 0.01% Tween 20. Reaction mixes were prepared by combining 10 μl cells at stated OD_600_ or spore density with recombinant Zt6, restrictocin (Sigma) or RNaseA (Qiagen) at 20 μM final concentration and distilled H_2_O added to a final reaction volume of 30 μl. Reactions were incubated for 24 h with gentle agitation. Serial dilutions of 10 μl aliquots of each reaction were prepared and plated on appropriate selection media and grown at 30°C (*S. cerevisiae*, *P. pastoris*, *E. coli*) or 16°C (*Z. tritici*) until colonies appeared.

## Results

### A *Z. tritici* gene *Zt6*, encoding a small secreted ribonuclease, shows biphasic upregulation during infection of susceptible wheat

A recent transcriptomic analysis of *Z. tritici* isolate IPO323 infection of the susceptible wheat cv. Riband (Rudd *et al.*, 2015) revealed *in planta* expression of a secreted ribonuclease (*Mycgr3G38105*, hereafter referred to as *Zt6*) of the N1/T1 class. Expression at all timepoints during wheat infection was higher than during growth in Czapek-Dox broth (CDB) (Fig. 1a). *Zt6* exhibited a double-peak expression pattern during wheat infection, with maximal expression at 1 dpi with a secondary peak at 14 dpi (Fig. 1a). These timepoints coincide with spore germination immediately following leaf surface inoculation (1 dpi) and the necrotrophic phase of the fungal life cycle (14 dpi), respectively. Expression of *Zt6* and several other genes encoding effector proteins (Kettles *et al.*, 2017) in an *in vitro* culture of *Z. tritici* IPO323 grown in nutrient-rich (YPD) medium vs *in planta*, at 1 dpi on susceptible wheat cv. Riband was also analysed by qRT-PCR (Fig. 1b). In agreement with our previous data (Rudd *et al.*, 2015), *Zt6* expression was considerably higher at 1 dpi during wheat infection compared to *in vitro* growth in YPD culture. Furthermore, induction of *Zt6* was more rapid than induction of several other *Z. tritici* candidate effectors tested (Fig. 1b). Collectively, these data indicate that *Zt6* is rapidly induced following spore germination on the wheat leaf surface in comparison to growth *in vitro*.

**Figure 1.**
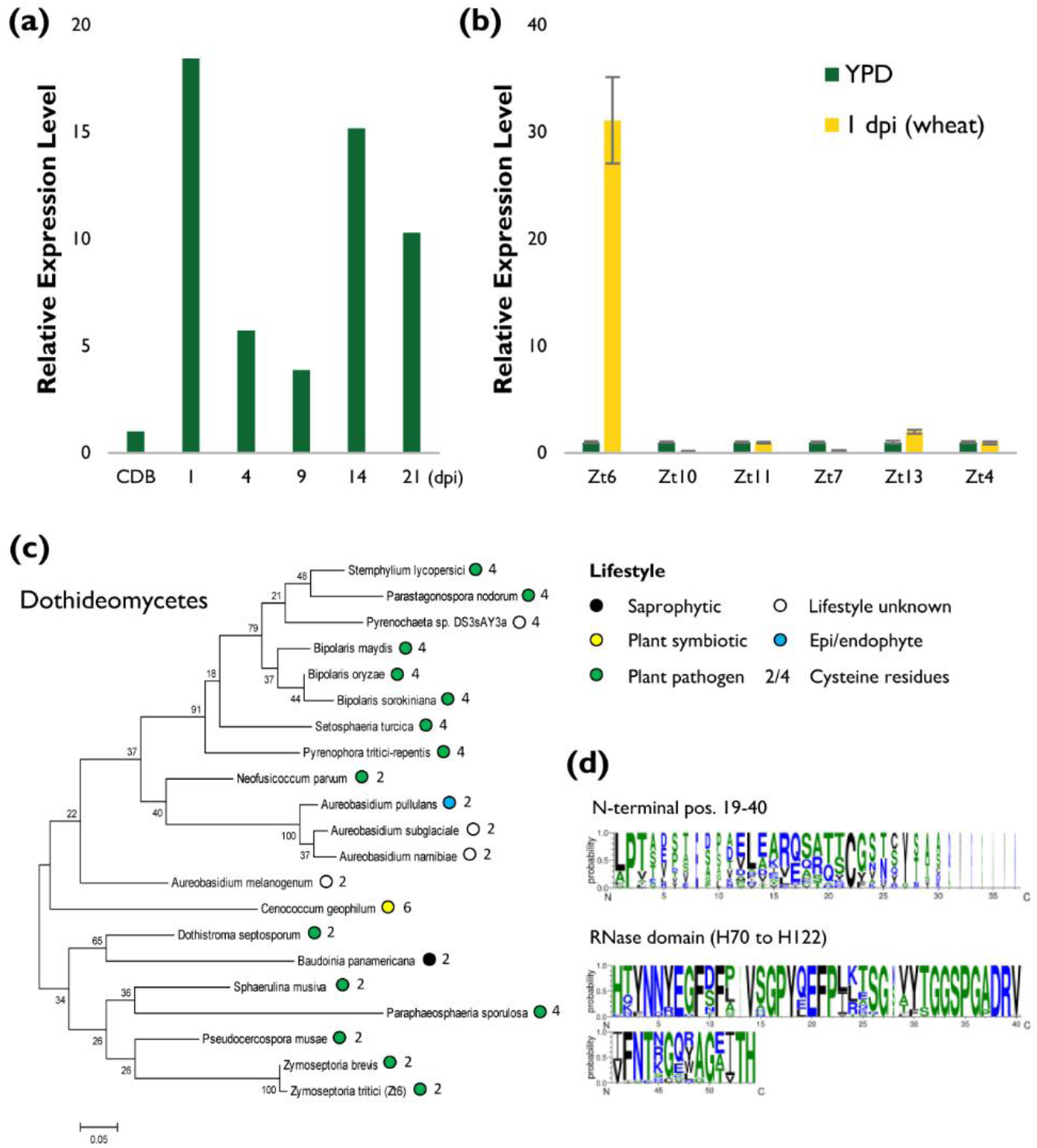
Transcriptomic and phylogenetic analysis of *Z. tritici* effector Zt6. (a) RNAseq expression profile of *Zt6* (*Mycgr3G38105*) from *Z. tritici* IPO323 infection timecourse (1-21 dpi) of susceptible wheat cv. Riband in comparison to growth in Czapek-Dox broth (CDB) (Rudd *et al.*, 2015). (b) qRT-PCR expression profile of *Zt6* from *Z. tritici* IPO323 grown in YPD broth in comparison to 1 dpi growth on susceptible wheat cv. Riband. (c) Phylogenetic representation of closest twenty Zt6 orthologs within the Dothideomycota. Coloured circles represent lifestyle of each species where orthologs are present. Sequences obtained by blastp (NCBI) and aligned using ClustalW. Tree constructed using Maximum Likelihood (ML) method with 100 bootstrap replicates (MEGA6). (d) WebLogo representations of the 22-amino acid N-terminal (residues 19-40) and RNase core (residues 70-122) domains of Zt6 and 20 closest Dothideomycete orthologs.

### *Zt6* is widely conserved within the Dothideomycetes

*Zt6* encodes a protein of 137 amino acids, with residues 1-18 corresponding to a secretion signal peptide (SP) predicted using SignalP and TargetP. The predicted RNase domain represents most of the mature protein sequence and is characterised by a histidine-glutamic acid-histidine (HEH) catalytic triad at amino acid positions 70, 88 and 122 respectively. Blastp analysis using the mature Zt6 protein sequence identified no paralogous genes in *Z. tritici* and revealed that homologs (between one and three per species) are present in many Ascomycete fungi, and in all species with sequenced genomes within the class Dothideomycetes (Fig. 1c). The closest homologs were found in species with a pathogenic lifestyle, although singular examples were also found in fungi described as epi/endophytic, saprophytic or symbiotic. Mature Zt6 protein (residues 19-137) contains two cysteine residues (residues 36, and 133) which are not predicted to form a disulphide bridge (DiANNA, SCRATCH protein predictor DIpro). This 2-cysteine form of secreted ribonuclease are also found in the majority of most closely related plant pathogenic fungal species including *Zymoseptoria brevis* and *Pseudocercospora musae which like Z. tritici are also members of the Mycosphaerellaceae* (Fig. 1c). However, many pathogens including important wheat leaf-infecting species such as *Parastagonospora nodorum, Bipolaris sorokiniana* and *Pyrenophora tritici-repentis* encode a 4-cysteine form which may impact protein folding. Analysis of overall sequence diversity between Zt6 homologues indicated that most variation is seen at the N-terminal end of the mature protein (residues 19-40 shown) whereas there is a high level of conservation within the RNase functional domain (H70 to H122 shown) (Fig. 1d).

### Zt6 encodes a functional cytotoxic protein active against wheat leaf cells

To assess potential Zt6 toxicity in the natural host (wheat), we used biolistic bombardment to coexpress a GFP transgene alongside mature Zt6 (-SP) (Fig. 2b, c) within host epidermal cells. In comparison to the GFP + EV control treatment, GFP + Zt6 (-SP) almost completely abolished GFP fluorescence, thus indicating potent cellular toxicity when expressed in wheat leaf cell cytoplasm. This data suggested that the mature Zt6 protein when expressed inside plant cells acted as a potent translational inhibitor. This data also established that the wheat bombardment assay was a suitable and high throughput system for the further characterisation of the Zt6 protein activity.

**Figure 2.**
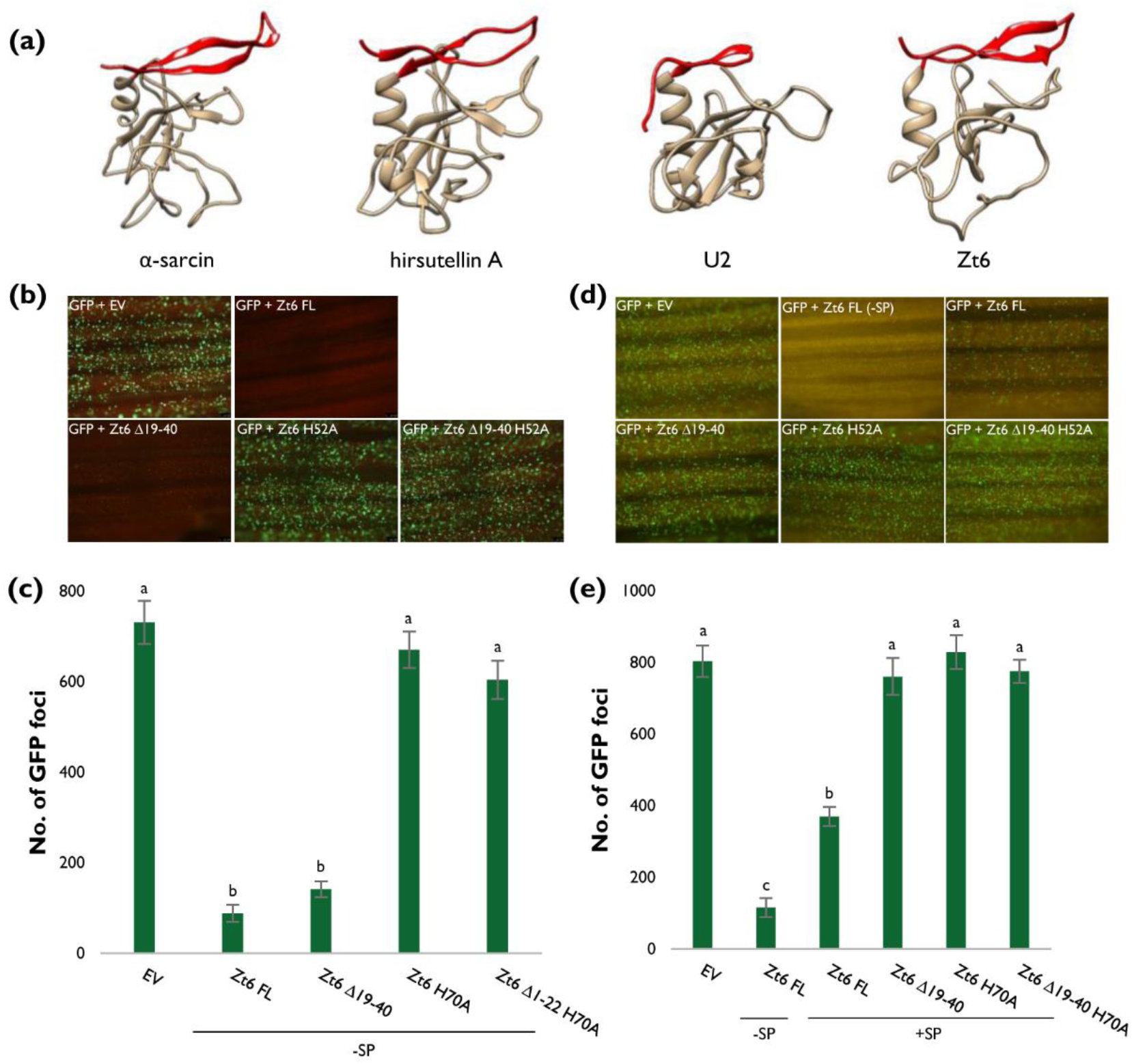
Zt6 inhibits transgene (GFP) expression in wheat leaves. (a) Protein folding prediction using Phyre2 for Zt6 alongside known structures of two ribotoxic RNases (α-sarcin, hirsutellin A) and the non-cytotoxic RNase U2. N-terminal loops (residues 19-40 for Zt6) highlighted in red. Images produced using Chimera Molecular Modeling System (UCSF). (b-e) Biolistic co-bombardment of GFP-and Zt6-expressing constructs (full-length (FL), N-terminal loop mutant (Δ19-40), catalytic mutant (H70A), double mutant (Δ19-40 + H70A) into wheat cv. Apogee leaves. Zt6 variants were expressed both without signal peptide (-SP) (b, c) and with signal peptide (+SP) (d, e). All leaves were photographed at 2 dpi and GFP foci counted using Metamorph software (Molecular Devices). Letters indicate differences at *P*<0.001 as determined by t-probabilities within a generalised linear model (GLM).

### The N-terminal loop region of the mature Zt6 RNase likely functions in enabling protein re-entry into wheat cells

It is known that the N-terminal region of secreted RNases can play important roles in cellular toxicity. The ribotoxin class of RNases for example have elongated N-terminal loops in a β-sheet-hinge-β-sheet structure that has been demonstrated to play roles in ribosome binding and postulated to be involved in cellular uptake (Garcia-Ortega *et al.*, 2002; García-Mayoral *et al.*, 2005), both important features that contribute to these molecules cytotoxicity. In contrast, non-toxic RNases have comparatively short N-terminal loops (Lacadena *et al.*, 2007). A structural prediction for Zt6 using Phyre2 (Kelley *et al.*, 2015) suggested an N-terminal loop region exists with similarity to the atypical ribotoxin hirsutellin A from the mite fungal pathogen *Hirsutella thompsonii* (Herrero-Galán et al., 2008) (Fig. 2a). We speculated that this region may be important for Zt6 function and that Zt6 may exhibit some characteristics of a ribotoxic RNase.

To investigate the contribution of the N-terminal loop to cytotoxicity, a *Zt6 Δ19-40* mutant missing the first 22 amino acids of the mature protein (Fig. 2a, shown in red) was co-expressed with GFP in the wheat biolistic system. This resulted in a near-complete absence of GFP fluorescence, indicating that this truncation mutant retains near wild-type activity when expressed intracellularly (ie without the secretion signal sequence). In contrast, co-expression of an RNase catalytic mutant (H70A) or the double mutant (Δ19-40, H70A) permitted GFP fluorescence at levels similar to those of the EV control (Fig. 2b, c) indicating a near-complete absence of toxicity. Together, this indicated that RNase activity is required for Zt6 toxicity but that an N-terminal loop is dispensable for this activity when the protein is retained (or expressed) intracellularly.

To test whether the N-terminal loop region might instead be important for protein uptake into wheat cells, a complementary set of bombardments were conducted with both mature Zt6 protein and the mutants described above, with a re-introduced secretion signal to direct the Zt6 protein to the apoplastic space of wheat cells and then monitor effects on intracellular GFP fluorescence (Fig. 2d, e). Full-length Zt6 (containing its secretion signal) was still able to strongly inhibit GFP fluorescence during co-bombardment, suggesting this secreted fungal protein can re-enter the host cell cytoplasm, where presumably it exerts its cytotoxic activity. Interestingly, and in stark contrast, Zt6 Δ19-40 expressed with the secretion signal induced no or very little host cell death as GFP expression was not compromised. In fact, the activity of the Δ19-40 mutant was approximately equivalent to the EV control and the Zt6 H70A catalytic mutant (Fig. 2d, e). This data indicates that the N-terminal 22 amino acids of mature Zt6 are important for toxicity against wheat cells only when the protein is initially directed to the apoplastic space of wheat leaves. The most likely explanation for this effect is that the N-terminal loop may be required for Zt6 re-entry into host cells.

### Both mature Zt6 and the Δ19-40 mutant protein are fully functional RNases active against rRNA

Cytotoxic RNases classed as ribotoxins, are known to exert cytotoxicity by disruption of ribosomal RNA (rRNA)-protein complexes or cleavage of rRNA with varying degrees of specificity. Despite numerous attempts, we were unable to successfully express FL recombinant Zt6 in *E. coli* or *P. pastoris* protein expression systems. We therefore utilised *in vitro* transcription coupled with a cell-free protein expression system to assess ribotoxin activity of mature Zt6 and its mutant versions Δ19-40, H70A and Δ19-40 + H70A towards native, functional ribosomes isolated from animal and plant cells (Fig. 3). Full-length Zt6 showed a ribotoxin-like activity cleaving both rabbit (Fig. 3a) and wheat (Fig. 3b) rRNA in an apparent semi-specific manner, yielding distinct cleavage fragments from both substrates, with greater degradation apparent from rabbit rRNA compared to wheat rRNA. However, whilst Zt6 H70A and Zt6 Δ19-40 + H70A displayed no catalytic activity against either substrate, Zt6 Δ19-40 activity was indistinguishable from that of wild-type Zt6 (Fig. 3a, b). Therefore, by contrast to the bona-fide ribotoxins, the N-terminal loop appears to be dispensable for Zt6 RNase activity towards native rRNA. To further assess the catalytic activity of Zt6 FL and Zt6 Δ19-40, assays were conducted against both rabbit and wheat rRNA with shortened reaction times (Fig. 3c, d). These experiments indicated that against rabbit rRNA, reactions containing either Zt6 FL or Zt6 Δ19-40 progressed to completion within 10 minutes. Longer incubation did not result in further rRNA degradation (Fig. 3c). In contrast, degradation of wheat rRNA was slower for both Zt6 FL and Zt6 Δ19-40 (Fig. 3d), with reactions progressing to completion between 30-60 minutes’ incubation. The fragments produced from degradation of wheat rRNA by Zt6 FL and Zt6 Δ19-40 were examined by Bioanalyzer (Fig. S1). This analysis confirmed that, similarly to ribotoxins, both Zt6 FL and Zt6 Δ19-40 primarily degrade (or attack) the 28S rRNA subunit in a cell-free expression system.

**Figure 3.**
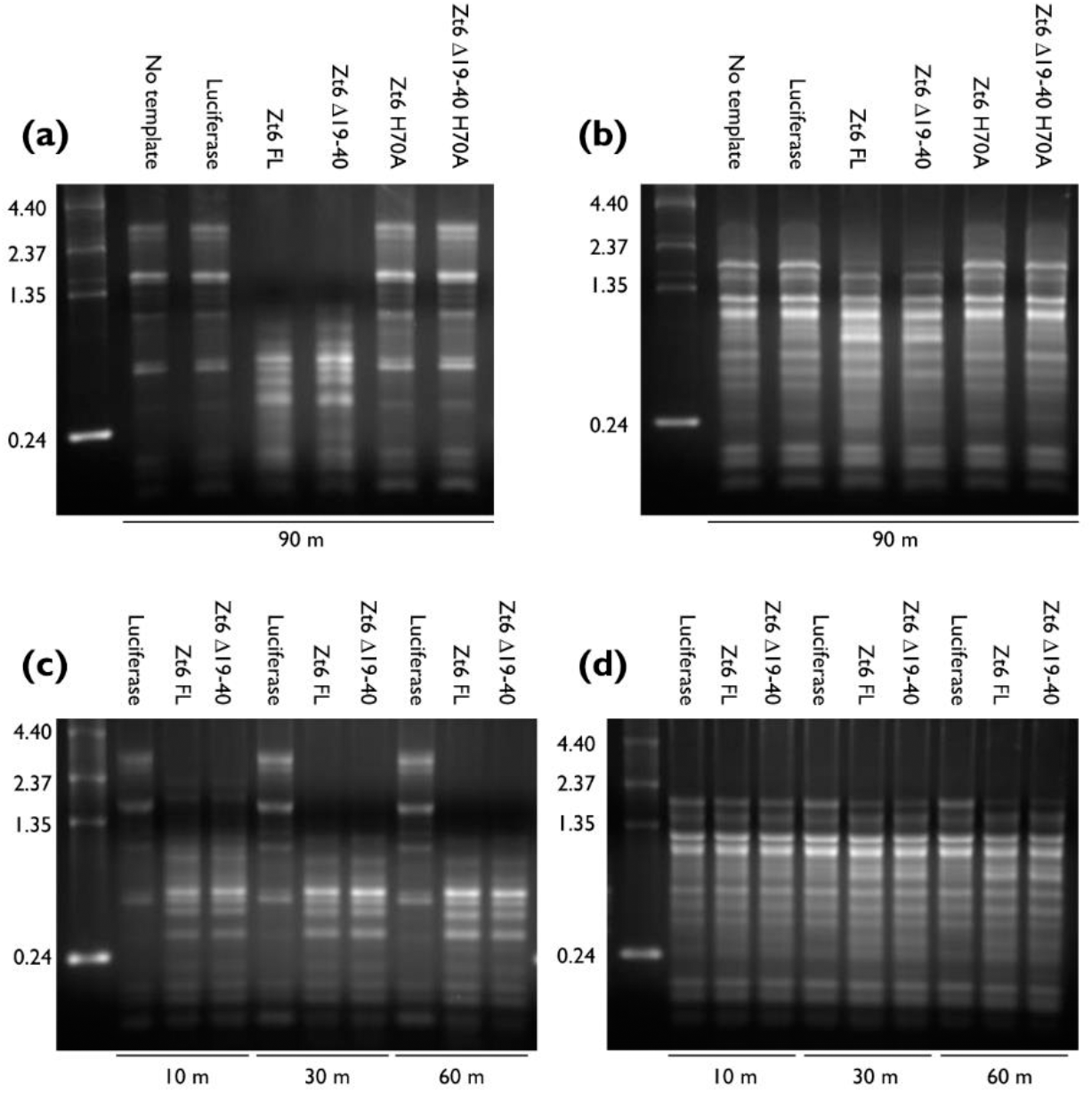
Zt6 is a functional RNase with catalytic activity against native rRNA from animal and plant ribosomes. Gel electrophoresis of recovered rRNA from rabbit reticulocyte lysate (a, c) and wheat germ extract (b, d) producing recombinant Zt6 FL, Zt6 Δ19-40, Zt6 H70A or Zt6 Δ19-40 H70A proteins by *in vitro* transcription and cell-free translation. Reactions were incubated for 90 min (a, b) and 10-60 min (c, d) prior to RNA recovery. No RNA template or Luciferase RNA template reactions as negative controls.

To further confirm this ribonuclease activity, we used a *P. pastoris* expression system to produce and purify a small quantity of Zt6 Δ19-40 protein. This was tested for RNase activity against both rabbit and wheat ribosomes alongside the highly-specific ribotoxic RNase restrictocin from *Aspergillus restrictus* and the non-specific RNase A from *Bos taurus* (Fig. S2). These experiments revealed that purified Zt6 Δ19-40 had activity similar to Zt6 Δ19-40 produced in the cell-free system. The activity of Zt6 Δ19-40 was dissimilar both to the activity of restrictocin which specifically generates the α-fragment from 28S rRNA, and RNase A which non-specifically degrades all RNA (Fig. S2). Zt6 Δ19-40 displayed non-specific RNase activity against denatured wheat total RNA, similar to RNase A (Fig. S3). Thus indicating that ribosomal structure confers specificity for Zt6 cleavage activity.

### The cytotoxic activity of Zt6 is conserved in the non-host model plant *N. benthamiana*

To assess whether Zt6 activity extended beyond the host range of *Z. tritici*, we used Agrobacterium-mediated transient expression (Agroexpression) to express Zt6 FL and mutant forms in the non-host tobacco *N. benthamiana* (Fig. 4). These experiments revealed that Zt6 is a highly potent inducer of cell death in this plant species with Zt6-induced symptoms first visible as early as at 2 dpi, which progressed to complete necrosis by 4 dpi (Fig. 4a). In comparison, symptoms induced by other previously characterised *Z. tritici* cell-death inducing effectors e.g. Zt9 (Kettles *et al.*, 2017) did not develop until 4 dpi, with widespread cell death in the infiltrated zone not clearly pronounced until 6 dpi (Fig. 4a). Furthermore, Zt6 induced rapid cell death irrespective of whether it was directed to the cytosol or the apoplast i.e. in the absence or the presence of its native SP, respectively (Fig. 4a). In contrast, the previously described Zt9 effector induced cell death only when directed to the apoplast, thus indicating that the mechanism upon which Zt6 induces cell death is distinct. Indeed, by contrast to several other previously characterised *Z. tritici* effectors (Kettles *et al.*, 2017) which induced light- and expression level-dependent cell death, Zt6 induced cell death in a light-and expression level-independent manner (Fig. 4b, c). Several *Z. tritici* effectors are also known to induce *N. benthamiana* cell death dependent on the Brassinosteroid Insensitive 1 (BRI1)-Associated Receptor Kinase 1 (BAK1) and Suppressor of BIR1-1 (SOBIR1) receptor-like kinases (RLKs). In contrast, we found that Zt6 induced cell death independently of these two RLKs (Fig. S4).

**Figure 4.**
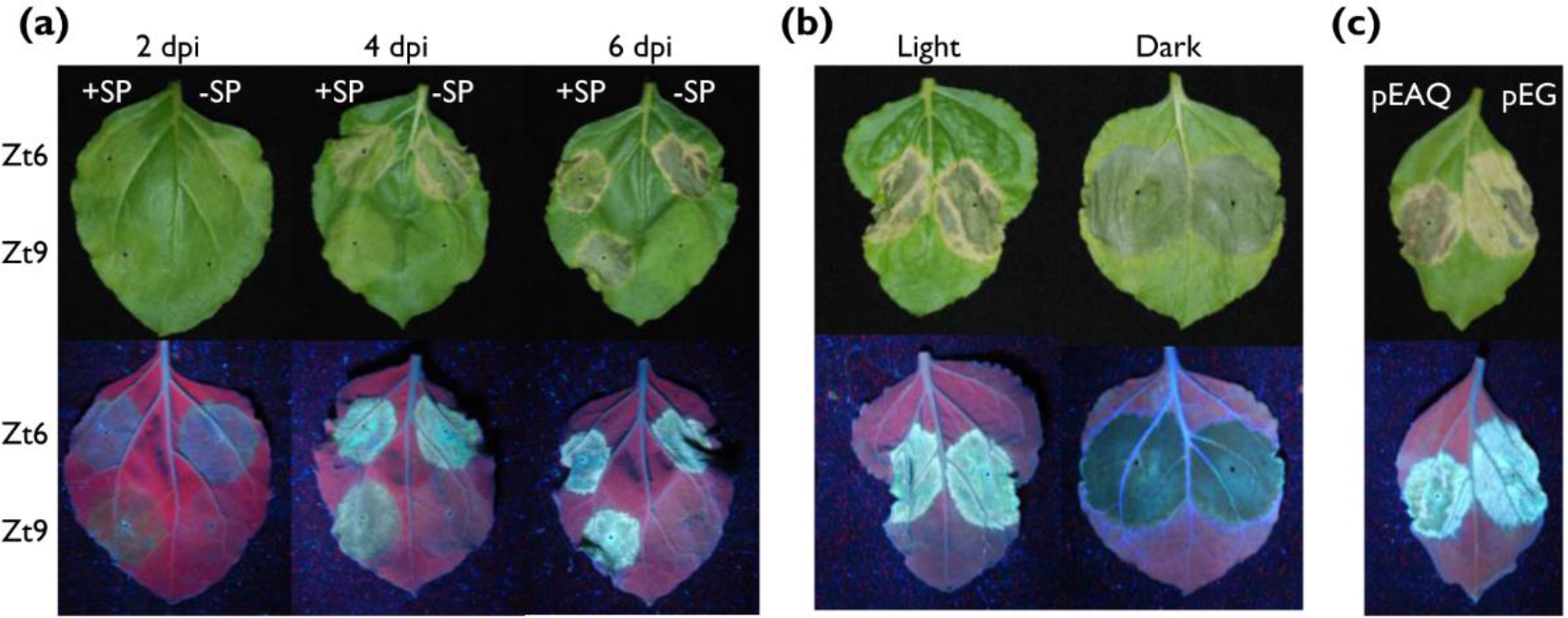
Zt6 is a potent inducer of cell death in the nonhost plant *N. benthamiana*. (a) Timecourse of Zt6-induced cell death in comparison with *Z. tritici* effector Zt9 (Kettles *et al.*, 2017) following Agroexpression. Leaves were photographed at 2, 4 and 6 days post inoculation (dpi) and both genes were expressed +/-their native secretion signal peptides (SP). (b) Zt6 (+SP) Agroexpressed in *N. benthamiana* leaves which were subsequently kept under a 16 h day-night cycle (Light) or in 24 h darkness (Dark) for 6 d. Leaves from the same plant are shown. (c) Zt6 (+SP) Agroexpressed from either the pEAQ-HT-DEST3 (pEAQ) or pEARLEYGATE101 (pEG) vector and leaves assessed at 7 dpi.

### Zt6 is toxic to prokaryotes and yeasts, but not to *Z. tritici*

To assess the contribution of Zt6 to *Z. tritici* virulence on wheat, we created targeted gene deletion mutants using Agrobacterium-mediated transformation. When tested in a wheat infection bioassay, three independent Δ*zt6* transformants displayed similar virulence as the background control strain (Δ*ku70*) on wheat cv. Riband (Fig. S5). That Zt6 appears dispensable for wheat infection, coupled with the unusual double-peak expression pattern during *in planta* infection (Fig. 1a, b) and potent cytotoxicity against plant cells (Fig. 2, 4) led us to speculate that Zt6 may have an alternate role in microbial competition and niche protection. That Zt6 might also exhibit broad antimicrobial toxicity was initially indicated by our failure to produce full length recombinant protein in all tested strains of *E. coli* and *P. pastoris*. All colonies recovered as transformants from either system failed to produce protein, and resequencing revealed they contained mutated, inactivated copies of Zt6 (data not shown). However, we were able to produce small amounts of the Zt6 Δ19-40 mutant in *P. pastoris*. This protein was therefore used in *in vitro* microbial toxicity assays alongside the cytotoxic ribotoxin restrictocin. Restrictocin exhibited potent toxicity against all microorganisms tested: the prokaryote *E. coli*, the yeasts *S. cerevisiae* and *P. pastoris*, and the filamentous fungus *Z. tritici* (Fig. 5a-d). In contrast, Zt6 Δ19-40 exhibited toxic activity against *E. coli*, *S. cerevisiae* and *P. pastoris* but had no activity against *Z. tritici*. Collectively, these results indicate that Zt6 is cytotoxic against bacteria and the lower yeasts but not towards *Z. tritici* itself.

**Figure 5.**
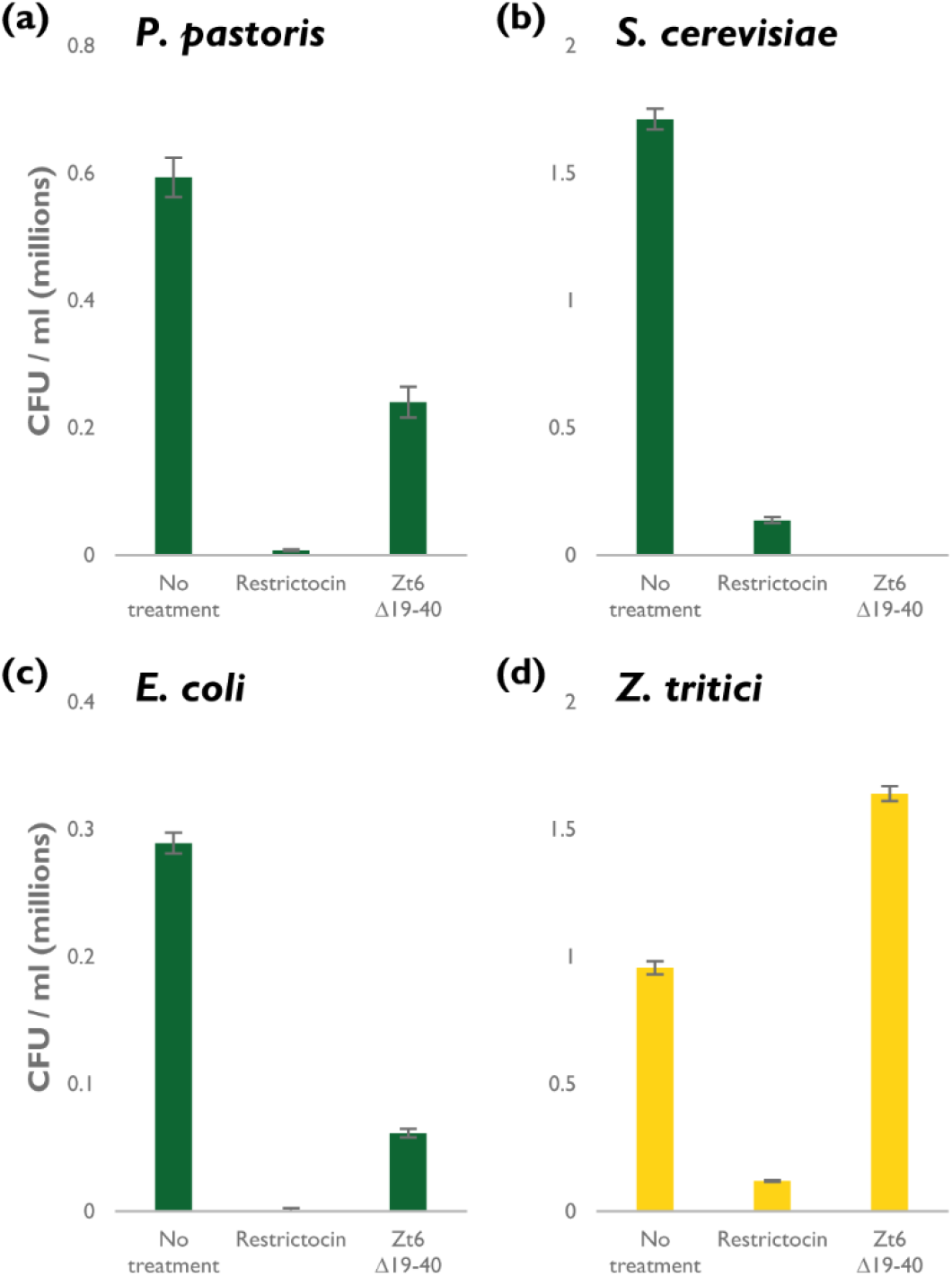
Zt6 is toxic to bacteria and yeasts but not *Z. tritici*. Recombinant Zt6 Δ19-40 and the ribotoxin restrictocin incubated with (a) *P. pastoris*, (b) *S. cerevisiae*, (c) *E. coli* and (d) *Z. tritici* at 20 μM concentration.

### Discussion

Here we describe the characterisation of Zt6, a small secreted ribonuclease with a ribotoxin-like activity from the wheat pathogen *Z. tritici* that exhibited potent cytotoxicity towards wheat, tobacco, bacterial and yeast cells. No other secreted protein from this pathogen has to date been demonstrated to have such broad-spectrum cytotoxicity. The MgNLP effector from *Z. tritici* induces rapid cell-death in dicotyledonous plants (*A. thaliana*, *N. benthamiana*) but does not have the same effect in wheat (Motteram *et al.*, 2009; Kettles *et al.*, 2017). Additionally, the secreted proteins ZtNIP1 and ZtNIP2 are reported to induce cultivar-specific necrosis/chlorosis in wheat (M’Barek *et al.*, 2015). We recently described a set of *Z. tritici* effectors that induce cell death in *N. benthamiana* (Kettles *et al.*, 2017). However, Zt6 has a cytotoxicity spectrum that goes beyond that demonstrated against plant cells in these previous studies to include yeasts and bacteria.

In our recent work, we demonstrated the cell death-inducing activity of 14 *Z. tritici* secreted proteins in the non-host plant *N. benthamiana*. Cell death induction by these effectors appears to be a result of an active recognition in the apoplast by cell surface immune receptors, as we showed it to be light-dependent and required the BAK1 and SOBIR1 RLKs. By contrast to that data, here we showed that Zt6-mediated cell death is both light-and BAK1/SOBIR1-independent (Fig. S4) and that Zt6 can induce cell death when directed either to the apoplast or the cytosol in both the non-host *N. benthamiana* (Fig. 4a, d, e) and a natural host plant, wheat (Fig. 2b-e). This distinguishes the functionality of Zt6 from all previously described effector candidates from *Z. tritici.*

The *Z. tritici* Δ*zt6* mutants displayed no loss of virulence during glasshouse infection assays on the susceptible wheat cultivar Riband (Fig. S5). A similar observation was made during infection of other susceptible wheat cultivars with the same strains (data not shown). This was surprising given the cytotoxic potency we demonstrated for Zt6. Given that Zt6 is the only ribonuclease identified in the *Z. tritici* secretome, it seems unlikely that functional redundancy with another secreted RNase accounts for absence of a loss-of-virulence phenotype. Though it is possible that other fungal proteins may also contribute to cell death induction perhaps using a distinct mechanism. However, from the data provided it is equally likely that the primary function of Zt6 may be to exert cytotoxic activity towards other microorganisms rather than host plant cells during natural *Z. tritici* infections.

Several studies (Mirzadi Gohari *et al.*, 2015; Rudd *et al.*, 2015; Palma-Guerrero *et al.*, 2016; Palma-Guerrero *et al.*, 2017) have described numerous putative *Z. tritici* effectors based on increased expression *in planta*. Particular focus has been made on those effectors with peak expression shortly before or during the fungal transition to necrotrophy. In contrast, Zt6 displays a rapid induction upon spore germination on the leaf surface at 1 dpi (Fig. 1a, b). This induction is more rapid and stronger than for several other previously characterised effectors (Fig. 1b). Given the high level of expression, the phase of early fungal growth on the leaf surface would represent the most likely time for Zt6-induced host cell death to occur. As it does not (Kema *et al.*, 1996) (personal observation) it suggests that Zt6 protein is unlikely to penetrate the waxy leaf cuticle to gain access to the exterior of leaf epidermal cells. This distinguishes Zt6 from, for example, the SnTox1 effector protein from *P. nodorum* which has recently been shown capable of direct penetration of the wheat leaf cuticle to elicit cell death in sensitive cultivars (Liu *et al.*, 2016). Therefore, at this point, we hypothesise that the function of Zt6 may be to clear the immediate area around the germinating spore of other potentially harmful or competing microorganisms. In contrast, the period of post-penetration asymptomatic colonisation represents the period of most likely Zt6 entry into leaf mesophyll cells. Given the reduction in Zt6 expression during this phase (4-9 dpi) (Fig. 1a), it suggests that Zt6 activity might be detrimental to fungal colonisation were it to induce significant host cell death during what is normally the asymptomatic phase of infection. Finally, the second peak of expression we have observed during the necrotrophic phase (14-21 dpi) (Fig. 1a), might be interpreted as the niche protective role of Zt6, minimising colonisation of the nutrient-rich necrotic tissue by opportunistic saprophytic microorganisms.

Structural predictions suggest that Zt6 displays some similarity to the ribotoxin hirsutellin A (Fig. 2a). Ribotoxins have long been known for their potent cytotoxicity and exquisite specificity for a cleavage of the phosphodiester backbone at a single nucleotide within the sarcin-ricin loop (SRL) of the 28S rRNA in eukaryotic ribosomes. Whilst Zt6 also displays considerable potency against numerous eukaryotic cells (Figs. 2, 4, 5), the mode of action appears different as Zt6 does not generate the characteristic α-fragment diagnostic of ribotoxin activity (Figs. 3, S2, S3). Whilst this distinction is clear, it is possible that Zt6 may share other features with ribotoxins in order to exert its cytotoxicity. In particular, this may relate to how Zt6 gains access to intracellular RNA when it is initially produced on the exterior of cells. This is exemplified by the requirement of a 22-amino acid N-terminal loop region for full potency. The classical ribotoxin α-sarcin also requires N-terminal amino acids 7-22 for full potency (Garcia-Ortega *et al.*, 2002). The α-sarcin Δ7-22 mutant retains ribonucleolytic activity against denatured RNA, but is unable to interact with intact ribosomes to generate the α-fragment. Additionally, the Δ7-22 mutant is less efficient at interacting with cell membranes thus impairing uptake by endocytosis (Garcia-Ortega *et al.*, 2002). It is speculated that other surface-exposed loops are important for full ribotoxin activity, either through interaction with lipid vesicles or the ribosome itself (Olombrada *et al.*, 2017). Given the similarity between Zt6 and α-sarcin in terms of requirement for a short N-terminal region for full potency, these other regions may also function in the ability of Zt6 to both gain access to cells and to exert ribonucleolytic activity.

Given the broad-spectrum toxicity of Zt6 against plant, yeast and bacterial cells, it is logical to assume there must exist some mechanism of self-protection in *Z. tritici*. In bacteria, there exist examples of bacteriocins displaying RNase activity (Riley & Wertz, 2002). Bacterial toxin-antitoxin systems are often based on toxic RNases, immunity to which is often conferred by the presence of dedicated immunity genes carried on the same plasmid (Cook *et al.*, 2013). The loss of membrane-localised susceptibility determinants in the producing cell can also confer RNase immunity (Riley & Wertz, 2002). Mammalian cells secrete RNases with antimicrobial function (Harder & Schröder, 2002; Pulido *et al.*, 2013) and are protected themselves by the well-characterised RNase Inhibitor (RI) protein (Dickson *et al.*, 2005). Human cells deficient in RI are themselves more vulnerable to their own secreted ribonucleases (Thomas *et al.*, 2016). It is therefore plausible that an analogous system exists to protect *Z. tritici*. However, the specific inhibitor involved is unknown as homologues of RI do not exist in the genomes of filamentous fungi. In addition, it is feasible that fundamental differences in lipid composition or physical structure between *Z. tritici* cell membranes and those of bacterial, yeast and plant cells does not permit Zt6 transit. As a phytopathogen, *Z. tritici* has also proven adept at the rapid evolution of resistance traits in response to the widespread application of chemical fungicides. Mechanisms of active efflux that confer improved fungicide tolerance may also offer a measure of self-protection to any toxic RNase that were able to re-enter the producing cell. However, at this point it remains completely unknown how filamentous fungi protect themselves against their own cytotoxic RNases, and this represents a fertile area for future study.

## Acknowledgements

We thank the Wheat Pathogenomics team (Rothamsted Research, UK) for fruitful discussions regarding this work and Rothamsted glasshouse staff for excellent plant care. The p75 *P. pastoris* expression vector and GS115 cells were kindly provided by Anna Avrova (James Hutton Institute, UK). Research was funded by the Institute Strategic Programme Grant “20:20 Wheat®” (grant no.BB/J/00426X/1) from the Biotechnology and Biological Sciences Research Council of the UK (BBSRC) and by the Rothamsted Institute Fellowship Programme.

## Author contributions

JR and KK initially conceived the project. GK, JR and KK designed the experiments. GK, CB, CS and GC conducted all experimental work, and GK analysed experimental data. GK, CS, JR and KK wrote the manuscript.

## References

Bolton MD, Van Esse HP, Vossen JH, De Jonge R, Stergiopoulos I, Stulemeijer IJE, Van Den Berg GCM, Borrás-Hidalgo O, Dekker HL, De Koster CG, et al. 2008. The novel Cladosporium fulvum lysin motif effector Ecp6 is a virulence factor with orthologues in other fungal species. Molecular Microbiology 69(1): 119–136.

Bowler J, Scott E, Tailor R, Scalliet G, Ray J, Csukai M. 2010. New capabilities for Mycosphaerella graminicola research. Molecular Plant Pathology 11(5): 691–704.

Cook GM, Robson JR, Frampton RA, McKenzie J, Przybilski R, Fineran PC, Arcus VL. 2013. Ribonucleases in bacterial toxin–antitoxin systems. Biochimica et Biophysica Acta (BBA) -Gene Regulatory Mechanisms 1829(6–7): 523–531.

de Jonge R, van Esse HP, Kombrink A, Shinya T, Desaki Y, Bours R, van der Krol S, Shibuya N, Joosten MHAJ, Thomma BPHJ. 2010. Conserved Fungal LysM Effector Ecp6 Prevents Chitin-Triggered Immunity in Plants. Science 329(5994): 953–955.

Dickson KA, Haigis MC, Raines RT. 2005. Ribonuclease Inhibitor: Structure and Function. Progress in nucleic acid research and molecular biology 80: 349–374.

Franco-Orozco B, Berepiki A, Ruiz O, Gamble L, Griffe LL, Wang S, Birch PRJ, Kanyuka K, Avrova A. 2017. A new proteinaceous pathogen-associated molecular pattern (PAMP) identified in Ascomycete fungi induces cell death in Solanaceae. New Phytologist.

García-Mayoral F, García-Ortega L, Álvarez-García E, Bruix M, Gavilanes JG, Martínez del Pozo A. 2005. Modeling the highly specific ribotoxin recognition of ribosomes. FEBS Letters 579(30): 6859–6864.

Garcia-Ortega L, Masip M, Mancheño JM, Oñaderra M, Lizarbe MA, Garcίa-Mayoral MF, Bruix M, del Pozo AM, Gavilanes JG. 2002. Deletion of the NH2-terminal ß-Hairpin of the Ribotoxin α-Sarcin Produces a Nontoxic but Active Ribonuclease. Journal of Biological Chemistry 277(21): 18632–18639.

Giraldo MC, Valent B. 2013. Filamentous plant pathogen effectors in action. Nat Rev Micro 11(11): 800–814.

Harder J, Schröder J. 2002. RNase 7, a Novel Innate Immune Defense Antimicrobial Protein of Healthy Human Skin. Journal of Biological Chemistry 277(48): 46779–46784.

Herrero-Galán E, Lacadena J, Martínez del Pozo A, Boucias DG, Olmo N, Oñaderra M, Gavilanes JG. 2008. The insecticidal protein hirsutellin A from the mite fungal pathogen Hirsutella thompsonii is a ribotoxin. Proteins: Structure, Function, and Bioinformatics 72(1): 217– 228.

Kao R, Martínez-Ruiz A, del Pozo AM, Crameri R, Davies J 2001. Mitogillin and Related Fungal Ribotoxins. In: Allen WN ed. Methods in Enzymology: Academic Press, 324–335.

Kelley LA, Mezulis S, Yates CM, Wass MN, Sternberg MJE. 2015. The Phyre2 web portal for protein modeling, prediction and analysis. Nat. Protocols 10(6): 845–858.

Kema GH, Yu D, Rijkenberg FHJ, Shaw MW, Baayen RP. 1996. Histology of the pathogenesis of Mycosphaerella graminicola in wheat. Phytopathology 86(7): 777–786.

Keon J, Antoniw J, Carzaniga R, Deller S, Ward JL, Baker JM, Beale MH, Hammond-Kosack K, Rudd JJ. 2007. Transcriptional Adaptation of Mycosphaerella graminicola to Programmed Cell Death (PCD) of Its Susceptible Wheat Host. Mol Plant Microbe Interact 20(2): 178–193.

Kettles G, Kanyuka K. 2016. Dissecting the molecular interactions between wheat and the fungal pathogen Zymoseptoria tritici. Frontiers in Plant Science 7.

Kettles GJ, Bayon C, Canning G, Rudd JJ, Kanyuka K. 2017. Apoplastic recognition of multiple candidate effectors from the wheat pathogen Zymoseptoria tritici in the nonhost plant Nicotiana benthamiana. New Phytologist 213(1): 338–350.

Lacadena J, Álvarez-García E, Carreras-Sangrà N, Herrero-Galán E, Alegre-Cebollada J, García-Ortega L, Oñaderra M, Gavilanes JG, Martínez del Pozo A. 2007. Fungal ribotoxins: molecular dissection of a family of natural killers. FEMS Microbiology Reviews 31(2): 212–237.

Lee WS, Rudd JJ, Hammond-Kosack KE, Kanyuka K. 2014. Mycosphaerella graminicola LysM effector-mediated stealth pathogenesis subverts recognition through both CERK1 and CEBiP homologues in wheat. Mol Plant Microbe Interact 27(3): 236–243.

Liu Z, Gao Y, Kim YM, Faris JD, Shelver WL, de Wit PJGM, Xu SS, Friesen TL. 2016. SnTox1, a Parastagonospora nodorum necrotrophic effector, is a dual-function protein that facilitates infection while protecting from wheat-produced chitinases. New Phytologist 211(3): 1052–1064.

Liu Z, Zhang Z, Faris JD, Oliver RP, Syme R, McDonald MC, McDonald BA, Solomon PS, Lu S, Shelver WL, et al. 2012. The cysteine rich necrotrophic effector SnTox1 produced by Stagonospora nodorum triggers susceptibility of wheat lines harboring Snn1. PLoS Pathog 8(1): e1002467.

M’Barek SB, Cordewener JH, Tabib Ghaffary SM, van der Lee TA, Liu Z, Mirzadi Gohari A, Mehrabi R, America AH, Robert O, Friesen TL, et al. 2015. FPLC and liquid-chromatography mass spectrometry identify candidate necrosis-inducing proteins from culture filtrates of the fungal wheat pathogen Zymoseptoria tritici. Fungal Genet Biol 79: 54–62.

Marshall R, Kombrink A, Motteram J, Loza-Reyes E, Lucas J, Hammond-Kosack KE, Thomma BP, Rudd JJ. 2011. Analysis of two in planta expressed LysM effector homologs from the fungus Mycosphaerella graminicola reveals novel functional properties and varying contributions to virulence on wheat. Plant Physiol 156(2): 756–769.

Mirzadi Gohari A, Ware SB, Wittenberg AH, Mehrabi R, Ben M'Barek S, Verstappen EC, van der Lee TA, Robert O, Schouten HJ, de Wit PP, et al. 2015. Effector discovery in the fungal wheat pathogen Zymoseptoria tritici. Mol Plant Pathol 16(9): 931–945.

Motteram J, Küfner I, Deller S, Brunner F, Hammond-Kosack KE, Nürnberger T, Rudd JJ. 2009. Molecular Characterization and Functional Analysis of MgNLP, the Sole NPP1 Domain–Containing Protein, from the Fungal Wheat Leaf Pathogen Mycosphaerella graminicola. Mol Plant Microbe Interact 22(7): 790–799.

Olombrada M, Lázaro-Gorines R, López-Rodríguez CJ, Martínez-del-Pozo A, Oñaderra M, Maestro-López M, Lacadena J, Gavilanes GJ, García-Ortega L. 2017. Fungal Ribotoxins: A Review of Potential Biotechnological Applications. Toxins 9(2).

Palma-Guerrero J, Ma X, Torriani SFF, Zala M, Francisco CS, Hartmann FE, Croll D, McDonald BA. 2017. Comparative Transcriptome Analyses in Zymoseptoria tritici Reveal Significant Differences in Gene Expression Among Strains During Plant Infection. Molecular Plant-Microbe Interactions 30(3): 231–244.

Palma-Guerrero J, Torriani SFF, Zala M, Carter D, Courbot M, Rudd JJ, McDonald BA, Croll D. 2016. Comparative transcriptomic analyses of Zymoseptoria tritici strains show complex lifestyle transitions and intraspecific variability in transcription profiles. Molecular Plant Pathology 17(6): 845–859.

Pedersen C, van Themaat EVL, McGuffin LJ, Abbott JC, Burgis TA, Barton G, Bindschedler LV, Lu X, Maekawa T, Weßling R, et al. 2012. Structure and evolution of barley powdery mildew effector candidates. BMC Genomics 13(1): 694.

Pliego C, Nowara D, Bonciani G, Gheorghe DM, Xu R, Surana P, Whigham E, Nettleton D, Bogdanove AJ, Wise RP, et al. 2013. Host-Induced Gene Silencing in Barley Powdery Mildew Reveals a Class of Ribonuclease-Like Effectors. Molecular Plant-Microbe Interactions 26(6): 633–642.

Pulido D, Torrent M, Andreu D, Nogués MV, Boix E. 2013. Two Human Host Defense Ribonucleases against Mycobacteria, the Eosinophil Cationic Protein (RNase 3) and RNase 7. Antimicrobial Agents and Chemotherapy 57(8): 3797–3805.

Riley MA, Wertz JE. 2002. Bacteriocins: Evolution, Ecology, and Application. Annual Review of Microbiology 56(1): 117–137.

Rovenich H, Boshoven JC, Thomma BPHJ. 2014. Filamentous pathogen effector functions: of pathogens, hosts and microbiomes. Current Opinion in Plant Biology 20: 96–103.

Rudd JJ, Kanyuka K, Hassani-Pak K, Derbyshire M, Andongabo A, Devonshire J, Lysenko A, Saqi M, Desai NM, Powers SJ, et al. 2015. Transcriptome and metabolite profiling of the infection cycle of Zymoseptoria tritici on wheat reveals a biphasic interaction with plant immunity involving differential pathogen chromosomal contributions and a variation on the hemibiotrophic lifestyle definition. Plant Physiol 167(3): 1158–1185.

Sánchez-Vallet A, Saleem-Batcha R, Kombrink A, Hansen G, Valkenburg DJ, Thomma BPHJ, Mesters JR. 2013. Fungal effector Ecp6 outcompetes host immune receptor for chitin binding through intrachain LysM dimerization. eLife 2: e00790.

Shi G, Zhang Z, Friesen TL, Raats D, Fahima T, Brueggeman RS, Lu S, Trick HN, Liu Z, Chao W, et al. 2016. The hijacking of a receptor kinase–driven pathway by a wheat fungal pathogen leads to disease. Science Advances 2(10).

Sparks CA, Jones HD 2014. Genetic Transformation of Wheat via Particle Bombardment. In: Henry RJ, Furtado A eds. Cereal Genomics: Methods and Protocols. Totowa, NJ: Humana Press, 201– 218.

Thomas SP, Kim E, Kim J, Raines RT. 2016. Knockout of the Ribonuclease Inhibitor Gene Leaves Human Cells Vulnerable to Secretory Ribonucleases. Biochemistry 55(46): 6359–6362.

Walsh MJ, Dodd JE, Hautbergue GM. 2013. Ribosome-inactivating proteins: Potent poisons and molecular tools. Virulence 4(8): 774–784.

Win J, Chaparro-Garcia A, Belhaj K, Saunders DGO, Yoshida K, Dong S, Schornack S, Zipfel C, Robatzek S, Hogenhout SA, et al. 2012. Effector Biology of Plant-Associated Organisms: Concepts and Perspectives. Cold Spring Harbor Symposia on Quantitative Biology 77: 235–247.

Zhong Z, Marcel TC, Hartmann FE, Ma X, Plissonneau C, Zala M, Ducasse A, Confais J, Compain J, Lapalu N, et al. 2017. A small secreted protein in Zymoseptoria tritici is responsible for avirulence on wheat cultivars carrying the Stb6 resistance gene. New Phytologist 214(2): 619–631.

